# BioThings SDK: a toolkit for building high-performance data APIs in biomedical research

**DOI:** 10.1101/2021.10.18.464256

**Authors:** Sebastien Lelong, Xinghua Zhou, Cyrus Afrasiabi, Zhongchao Qian, Marco Alvarado Cano, Ginger Tsueng, Jiwen Xin, Julia Mullen, Yao Yao, Ricardo Avila, Greg Taylor, Andrew I Su, Chunlei Wu

## Abstract

**Summary:** To meet the increased need of making biomedical resources more accessible and reusable, Web APIs or web services have become a common way to disseminate knowledge sources. The BioThings APIs are a collection of high-performance, scalable, annotation as a service APIs that automate the integration of biological annotations from disparate data sources. This collection of APIs currently includes MyGene.info, MyVariant.info, and MyChem.info for integrating annotations on genes, variants, and chemical compounds, respectively. These APIs are used by both individual researchers and application developers to simplify the process of annotation retrieval and identifier mapping. Here, we describe the BioThings Software Development Kit (SDK), a generalizable and reusable toolkit for integrating data from multiple disparate data sources and creating high-performance APIs. This toolkit allows users to easily create their own BioThings APIs for any data type of interest to them, as well as keep APIs up-to-date with their underlying data sources.

**Availability and implementation:** The BioThings SDK is built in Python and released via PyPI (https://pypi.org/project/biothings/).

Its source code is hosted at its github repository (https://github.com/biothings/biothings.api).

## Introduction

Concerns for reproducibility and the ethical considerations in ownership of publicly funded research has led to an increased effort to make biomedical research papers and data more accessible. While the open access/open data movement made biomedical research data more accessible, it did not enhance its re-usability. The FAIR data guiding principles were formed to address some of the issues and limitations of the open data movement (Wilkinson MD et al, 2016). For data to be both accessible, interoperable, and reusable, metadata standardization is needed. General online repositories like Figshare or Zenodo impose basic metadata requirements which help to standardize some basic metadata, but a lot of biomedical databases and repositories utilize their own metadata annotations (Singh, 2011)( European Organization For Nuclear Research, 2013). In addition to existing databases and repositories, web-based Application Programming Interfaces (APIs) play an important role in making biomedical research data more accessible and reusable and often have their own fragmented metadata.

We previously created high-performance, scalable, annotation as a service APIs that aggregated and provided gene and variant annotation data (Xin et al., 2016): MyGene.info (Xin et al., 2010) and MyVariant.info (Xin et al., 2014). These APIs aggregate identifiers and other metadata from gene-specific or variant-specific resources and serve up these annotations as a RESTful service effectively increasing the interoperability of data from various gene and variant specific resources. This allows users to address two key issues in bioinformatics pipelines that make keeping data up-to-date difficult: 1-data storage, 2-data fragmentation (ie-downloading, tracking, and updating data, vs constantly updating parsers for multiple resources). Our team has since built similar APIs for chemical entities (MyChem.info (Lelong et al., 2015)), disease entities (MyDisease.info (Xin et al., 2015)), and taxonomy (t.biothings.io/ (Lelong et al., 2017)). These APIs are collectively referred to as the **BioThings APIs**. Calling these BioThings APIs enable users to quickly perform id conversion and knowledge retrieval for ease of querying the respective data resources. By relying on Annotation as a service APIs, users reduce the need to download and store data from multiple resources, track and update said data locally, and resolve changes from multiple resources. This enhances the accessibility, interoperability, and reusability of data scattered across multiple resources.

The BioThings APIs were built with architecture demonstrated to be flexible and scalable. We currently have APIs that cover five different types of biomedical data out of the potentially hundreds if not thousands of biomedical entity types. To increase the FAIRness of different biomedical entity types, we encourage and empower other research groups to create and share their own APIs using the BioThings architecture. We reviewed the process of creating BioThings API, compared the architecture and applied the best practices and lessons learned from the process to create a python-based Software Development Kit (SDK). The SDK allows users to create a high-performance API once the user provides a data source parser. Once the API is created, the BioThings API client can be adapted for use with the API. Additionally, the BioThings SDK includes a data source management console which helps to handle data source monitoring and download scheduling (tasks important for keeping aggregated data up-to-date). Overall, we provide BioThings SDK as a system to rapidly increase the availability of high-quality, sustainable production-ready APIs to the research community.

## Implementation

The SDK consists of a data hub component and a web API component (**Figure 1**). The data hub processes each data source (“dump”) and stores the parsed JSON objects (“upload”) in MongoDB (MongoDB, 2021). All JSON objects can then be merged across data sources (based on the same primary keys) and indexed in Elasticsearch (Tong Z, 2013). The web API handles the user queries and returns back the query results. The API is built with the Python-based Tornado web framework (Tornadoweb.org, 2021), which is non-blocking and suitable for implementing high-performance APIs. The data hub also includes a set of utilities to monitor the data source files and triggers the automated data-downloads data-parsing and releasing. This feature greatly simplifies the data-processing burden and keeps the data always up-to-date. The SDK is regularly released as a package via PyPI (https://pypi.orq/project/biothings/) and its source code is hosted at the github repository (https://github.com/biothingsbiothings.api). Detailed documentation for the BioThings SDK (https://docs.biothings.io), a full tutorial, and preconfigured Dockerized development environment and a web interface (called “BioThings Studio”) are also provided (https://docs.biothings.io/en/latest/doc/studio.html).

**Figure 1.**
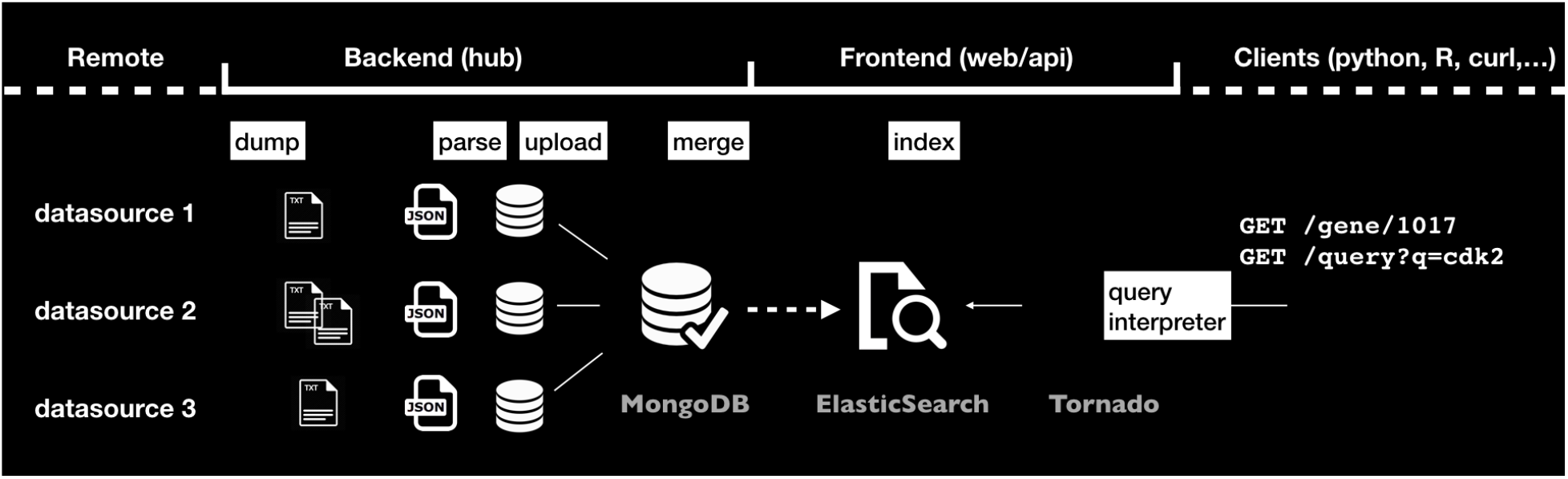
The overall architecture of the BioThings SDK. Its data hub (backend) component handles data source monitoring, parsing/uploading and then merging across all data sources. Its web API (frontend) component handles data indexing and processes user queries.

## Results

To create a RESTful API out of a data source (or add a data source into an existing API), a “data plugin” must be written. The data plugin will need to follow a simple but specific architecture consisting of three primary python modules: the *dumper.py*, which tells system where, how, and even how frequently to get the data; the *parser.py*, which tells the system how to parse and transform the data; and the *upload.py*, which tells the system how to load and merge the data into MongoDB. The BioThings SDK comes packaged with many helper functions which may be called in the data plugin files to reduce the amount of coding needed for common or redundant actions. For example, the SDK has helpers for downloading data via ftp, http, git, google drive, and more that can be used in the data plugin dumper script. In most straightforward scenarios, a data plugin can include just a *parser.py* and a *manifest.json* file with sufficient metadata (Lelong, 2020a), however, with dedicated *dumper.py* and *upload.py,* users have additional fine-tuned controls to customize the desired behavior (Lelong, 2020b). Once the data plugin has been created, it can be registered within the dashboard.

Registering the plugin within the dashboard allows the operator to manually trigger or schedule data updates and monitor the plugin for issues with uploading, merging, and indexing (Supplemental Figure 1). If the SDK is used to create an API for a single data source, each upload for the data into MongoDB can be considered a single build and can be configured as such. If there are multiple data sources, the build can be configured to handle the upload and merging of the data sources. For example, if a BioThings API like MyGene.info consisted of only a single data source (NCBI Gene), then each upload from the data plugin would have only the annotations available from NCBI entrez and each build would consist only of updates from NCBI Gene. In actuality MyGene.info provides gene-specific annotations from NCBI Gene, Ensembl, Uniprot, NetAffx, PharmGKB, UCSC, CPDB, ClinGen, REACTOME, and more. Since each data source has a different frequency with which updates are released, the data plugins are scheduled to update according to the releases. Any update will trigger a new merge and build with each JSON object in the database containing annotation data from all relevant data sources. Build configurations can be created and edited directly from the dashboard (Supplemental Figure 2).

Once the build has been completed, it will be sent to Elasticsearch for indexing. Elasticsearch can be customized to map/index specific fields/properties and this customization allows for the creation of customized API queries. Fields or properties not indexed by Elasticsearch will still exist within the data but cannot be queried. The SDK dashboard also has analysis tools for inspecting the data to automatically generate potential Elasticsearch. After the build is indexed by Elasticsearch, a new API can be created from within the dashboard (Supplemental Figure 3).

## Conclusions

We have created an SDK for building RESTful APIs for the biomedical research community. The BioThings SDK has been used to build over 50 APIs which collectively includes and exposes over 1.7 billion records. Though the usage of these APIs remains largely untracked, the monthly requests for the 5 APIs where usage was tracked exceeded 53 million (Supplemental Table 1). The BioThings SDK comes with a dashboard that allows the operator to easily manage some of the more redundant data wrangling tasks.

