## Supplemental Figures for "BioThings SDK: a toolkit for building high-performance data APIs in biomedical research"

Supplemental Figure 1: Once registered, data from a source (a data plugin) can be monitored and updated from within the dashboard

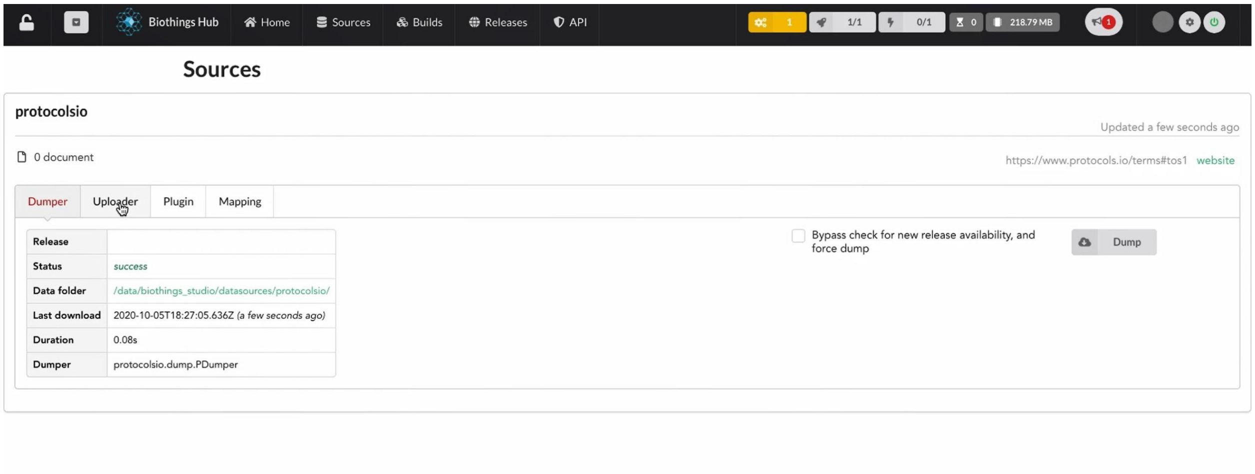


Supplemental Figure 2 - Build configurations can be created and edited directly from the dashboard

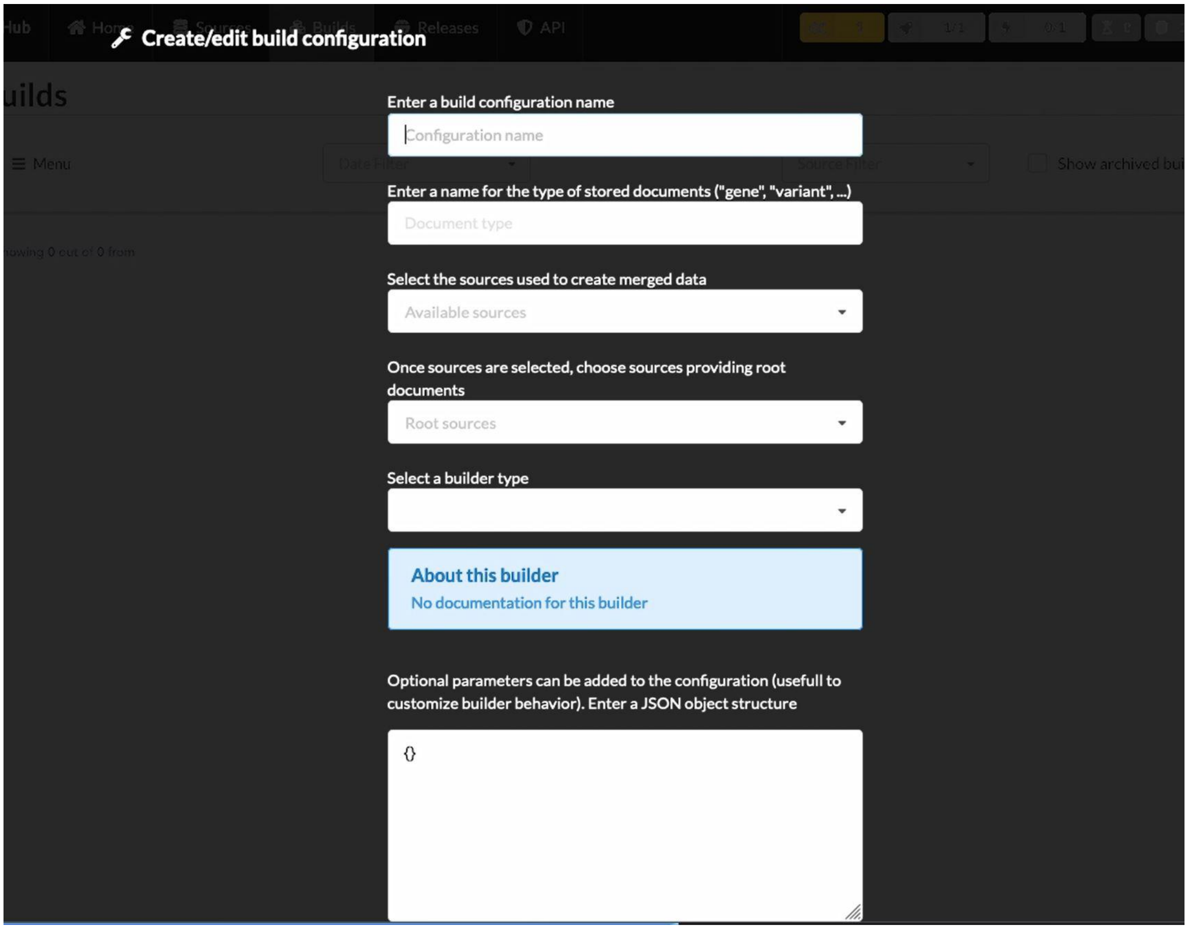

Supplemental Figure 3 - After the build is indexed by Elasticsearch, a new API can be created from within the dashboard

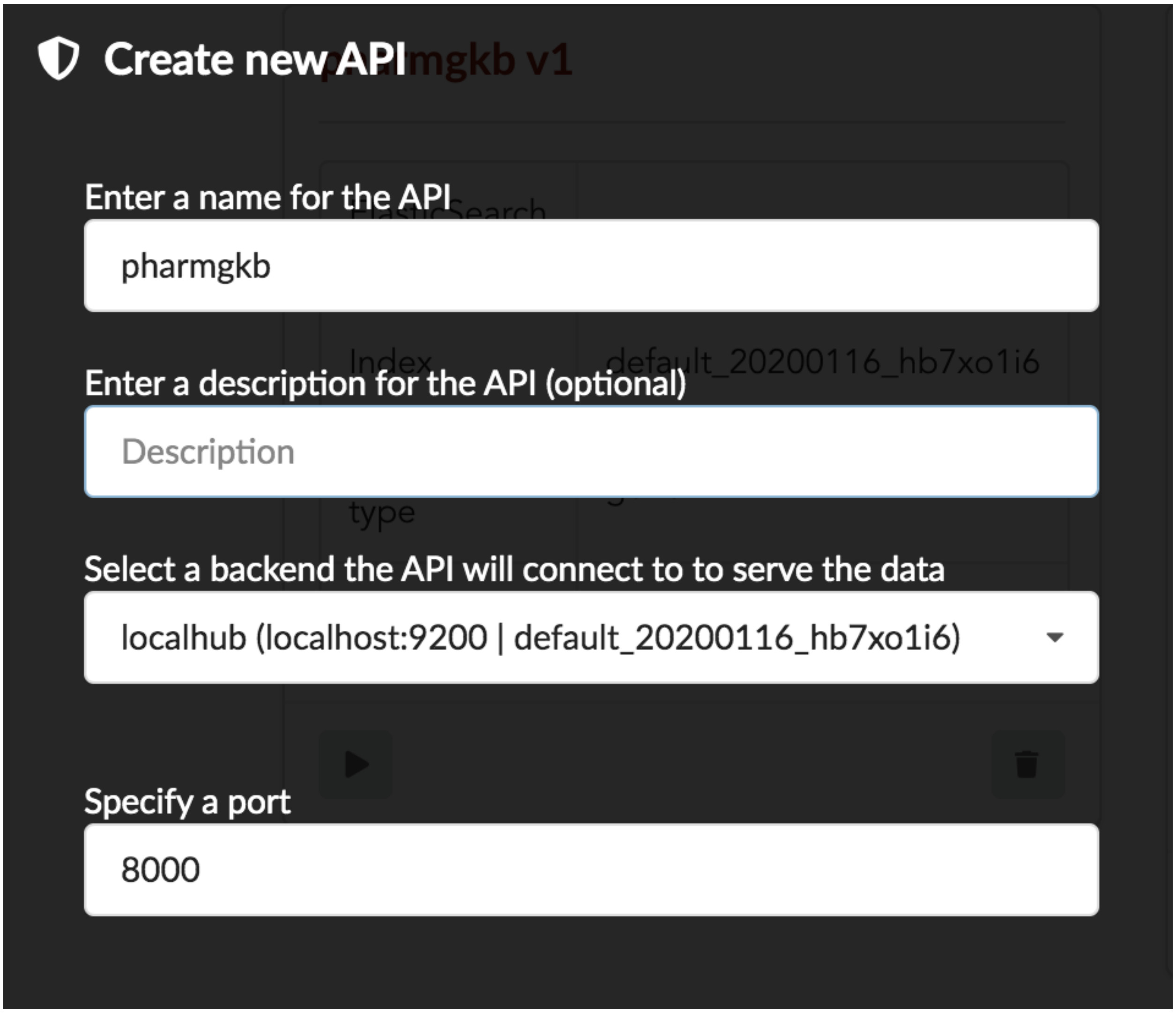
