## Supplemental Table 1 for "BioThings SDK: a toolkit for building high-performance data APIs in biomedical research"

| APIs | records | resource type | Requests within last 30 days of 10/15/2021 |
|---|---|---|---|
| MyGene.info | 46107797 | gene | 18,007,523 |
| MyVariant.info | 803819895 | variant | 27,177,600 |
| MyChem.info | 177383131 | chemcial | Not available |
| MyDisease.info | 126807 | disease | 962 |
| t.biothings.io | 126807 | taxonomy | 103 |
| Outbreak.info | 2,941,798 | COVID-19 Resources metadata, epidemiology, and genomics data | 8,688,332 |
| CViSB data API | Not available | Ebola, Lassa, COVID-19 data | Not available |
| N3C data API | Not available | COVID-19 data | Not available |
| NIAID data API | Not available | Allergy and Infectious Disease Resources metadata | Not available |
| **cord_cc** | 787 | geneset | Not available |
| **cord_genomic_entity** | 632 | genomicentity | Not available |
| **cord_cell** | 576 | cell | Not available |
| **cord_ma** | 1,594 | geneset | Not available |
| **agr** | 37,994 | association | Not available |
| **cord_protein** | 10,321 | protein | Not available |
| **cord_chemical** | 6,231 | chemical | Not available |
| **denovodb** | 271,069 | variant | Not available |
| **biggim** | 303,581,994 | association | Not available |
| **ccle** | 1,045,196 | variant | Not available |
| **clinical_risk_kp** | 471,872 | association | Not available |
| **cell_ontology** | 2,234 | cell | Not available |
| **cord_anatomy** | 2,730 | anatomy | Not available |
| **biomuta** | 3,737,897 | variant | Not available |
| **cord_bp** | 2,960 | geneset | Not available |
| **cord_disease** | 2,631 | disease | Not available |
| **dgidb** | 70,248 | association | Not available |
| **cord_gene** | 10,292 | gene | Not available |
| **DISEASES** | 5,703 | disease | Not available |
| **drug_response_kp** | 4,803,253 | association | Not available |
| **ebigene2phenotype** | 2,038 | gene | Not available |
| **geneset** | 58,745 | geneset | Not available |
| **go_bp** | 28,927 | geneset | Not available |
| **go_cc** | 4,189 | geneset | Not available |
| **go_mf** | 11,146 | geneset | Not available |
| **gwascatalog** | 87,941 | variant | Not available |
| **hpo** | 15,247 | phenotype | Not available |
| **idisk** | 919 | umls | Not available |
| **kaviar** | 213,390,302 | variant | Not available |
| **mgigene2phenotype** | 66,089 | gene | Not available |

| | | | |
|---|---|---|---|
| mrcoc | 46,064,731 | cooccurrence | Not available |
| multiomics_wellness_kp | 288,526 | association | Not available |
| pfocr | 55,309 | geneset | Not available |
| phenotype | 15,247 | phenotype | Not available |
| phewas | 3,144 | variant | Not available |
| pseudocap_go | 1,537 | gene | Not available |
| repodb | 2,211 | chemical | Not available |
| semmed | 36,248 | disease | Not available |
| semmed_anatomy | 2,750 | anatomy | Not available |
| semmedbp | 6,887 | geneset | Not available |
| semmedchemical | 58,812 | chemical | Not available |
| semmeddb | 114,383,742 | association | Not available |
| semmedgene | 33,339 | gene | Not available |
| semmedphenotype | 393 | phenotype | Not available |
| tcga_mut_freq_kp | 332,347 | association | Not available |
| text_mining_co_occurrence_kp | 21,000,000 | association | Not available |
| text_mining_targeted_association | 1,155,272 | association | Not available |
| textminingkp | 34 | protein | Not available |
| uberon | 13,898 | anatomy | Not available |
| umlschem | 58,812 | chemical | Not available |
| upheno_ontology | 13,600 | phenotype | Not available |
| Summary: | | | |
| Total APIs: | 60 | | |
| Total available records: | 1,741,764,831 | | |
| Total available monthly requests: | 53,874,520 | | |
